# Serotonin depletion impairs both Pavlovian and instrumental reversal learning in healthy humans

**DOI:** 10.1101/2020.04.26.062463

**Authors:** Jonathan W. Kanen, Annemieke M. Apergis-Schoute, Robyn Yellowlees, Frederique E. Arntz, Febe E. van der Flier, Annabel Price, Rudolf N. Cardinal, David M. Christmas, Luke Clark, Barbara J. Sahakian, Molly J. Crockett, Trevor W. Robbins

## Abstract

Serotonin is implicated in aversive processing and updating responses to changing environmental circumstances. Optimising behaviour to maximise reward and minimise punishment may require shifting strategies upon encountering new situations. Likewise, emotional reactions to threats are critical for survival yet must be modified as danger shifts from one source to another. Whilst numerous psychiatric disorders are characterised by behavioural and emotional inflexibility, few studies have examined the contribution of serotonin in humans. We modelled both processes in two independent experiments (N = 97), using instrumental and aversive Pavlovian reversal learning paradigms, respectively. Upon depleting the serotonin precursor tryptophan – in a double-blind randomised placebo-controlled design – healthy volunteers showed impairments in updating both behaviour and emotion to reflect changing contingencies. Reversal deficits in each domain, furthermore, were correlated with the extent of tryptophan depletion. These results translate findings in experimental animals to humans and have implications for the neurochemical basis of cognitive inflexibility.

## Introduction

Serotonin (5-HT; 5-hydroxytryptamine) is implicated in processing negative events and adapting previously learned responses to reflect new environmental circumstances (Cools et al. 2008). Behaviour that had been optimal for maximising reward and minimising punishment must at times be adjusted to navigate the world effectively. Likewise, reacting emotionally to threats is critical for survival yet must be modified as danger shifts from one source to another. Indeed, impairments in behavioural and emotional flexibility are pervasive in psychiatric disorders (Apergis-Schoute et al. 2017; APA 2013; Homan et al. 2019; Remijnse et al. 2006; Robbins et al. 2019; Waltz et al. 2007) and serotonergic dysfunction is widely implicated across diagnostic categories (Cunningham et al. 2020; Dayan and Huys 2009; Dersken et al. 2020; Phillips and Robbins 2020; Quednow et al. 2020; Zangrossi et al. 2020). Behavioural inflexibility is a hallmark of obsessive-compulsive disorder (OCD) and substance use disorders, for instance, where learned behaviours persist inappropriately despite adverse consequences (APA 2013). Aberrations in threat and safety learning are characteristic of post-traumatic stress disorder (PTSD; Homan et al., 2019; Milad et al., 2009) and other anxiety disorders (Kim et al., 2011; Marin et al., 2017), and are also a feature of OCD (Milad et al., 2013; Apergis-Schoute et al., 2017) and schizophrenia (Holt et al., 2012). Drugs thought to boost serotonin transmission – selective serotonin reuptake inhibitors (SSRIs) – are first line treatments for OCD (Fineberg et al. 2013) and PTSD (Baldwin et al. 2014). Schizophrenia, in which behavioural inflexibility has additionally been documented (Waltz et al. 2007), is treated with drugs that modulate serotonin in addition to dopamine, such as risperidone, a non-selective serotonin 2A (5-HT2A) receptor antagonist (Stahl, 2013). Drugs of this class can also be used to augment SSRI therapy in OCD (Fineberg et al. 2013).

Despite broad clinical relevance, the preponderance of evidence on how serotonin impacts behavioural adaptation comes from studies of non-human animals (e.g. Bari et al. 2010; Barlow et al. 2015; Clarke et al. 2004, 2007; Lapiz-Bluhm et al. 2009; Matias et al. 2017), whilst the role of serotonin in human threat and safety learning has received surprisingly little attention (Bauer 2015). Here, we studied healthy human volunteers to examine the effects of lowering serotonin on both behavioural and emotional flexibility in two independent experiments employing acute tryptophan depletion (ATD). Depleting tryptophan, serotonin’s biosynthetic precursor, decreases serotonin function (Bel & Artigas 1996; Bell et al. 2005; Biggio et al. 1974; Crockett et al. 2012a; Nishizawa et al. 1997). Reversal learning is a key laboratory assay of cognitive inflexibility that has been translated across species (Cools et al. 2008). Experiment 1 tested instrumental reversal learning. Individuals acquired an adaptive behaviour through trial and error learning (stimulus-response-outcome), and the correct response subsequently changed multiple times, necessitating cessation of the previous action and performing a new behaviour. Failure to adapt to new contingences is referred to as perseveration. Experiment 2 examined reversal learning in the Pavlovian domain (Apergis-Schoute et al. 2017; Schiller et al. 2008). Participants were presented with two cues (threatening faces, i.e. signs of aggressive conspecifics), one of which was sometimes paired with an electric shock, while the other was not (stimulus-outcome). A reversal phase followed, whereby the originally conditioned face became safe, and the initially safe stimulus was newly paired with shock. Under normal circumstances, anticipatory sympathetic nervous system arousal responses, manifested in perspiration, should track the presently threatening cue (Schiller et al. 2008).

Impairments in human instrumental reversal learning following ATD have been difficult to detect to date (Evers et al. 2005; Finger et al. 2007; Murphy et al. 2002; Park et al. 1994; Rogers et al. 1999; Talbot et al. 2006). Behaviour in previous studies, however, has not been reinforced with motivationally salient feedback. Consequently, there may not have been sufficient incentive to update or restrain action: any requirement for serotonin signalling to perform the task at hand may have been minimal enough to be unaffected by ATD (Faulkner and Deakin 2014). Indeed, the depletion achieved by ATD is relatively mild in comparison with the profound depletion that is possible in experimental animals (Bari et al. 2010; Clarke et al. 2004; 2007). Given the importance of serotonin in processing both aversive (Crockett et al. 2012b; Deakin 2013) and rewarding (Cohen et al. 2015; Matias et al. 2017; Seymour et al. 2012) outcomes, we used an innovative task (Figure 1) incorporating feedback that was markedly more salient than was used in previous reversal tasks (Evers et al. 2005; Finger et al. 2007; Murphy et al. 2002; Park et al. 1994; Rogers et al. 1999; Talbot et al. 2006). Prior studies that did not find a perseverative deficit following ATD employed largely probabilistic feedback (Evers et al. 2005; Finger et al. 2007; Murphy et al. 2002) and a single reversal (Finger et al. 2007; Murphy et al. 2002), or were used primarily to test higher order cognitive flexibility in the form of attentional set-shifting (Park et al. 1994; Rogers et al. 1999; Talbot et al. 2006). Meanwhile, evidence of a perseverative deficit following neurotoxic serotonin depletion, in the marmoset monkey orbitofrontal cortex (OFC), comes from a paradigm more similar to that employed in the present study: Clarke et al. (2004) used serial reversals on a deterministic schedule, and found a reversal deficit that emerged only beginning in the second reversal. We were therefore particularly interested in whether focusing on a later reversal phase may be key to uncovering perseveration following ATD in humans.

**Figure 1.**
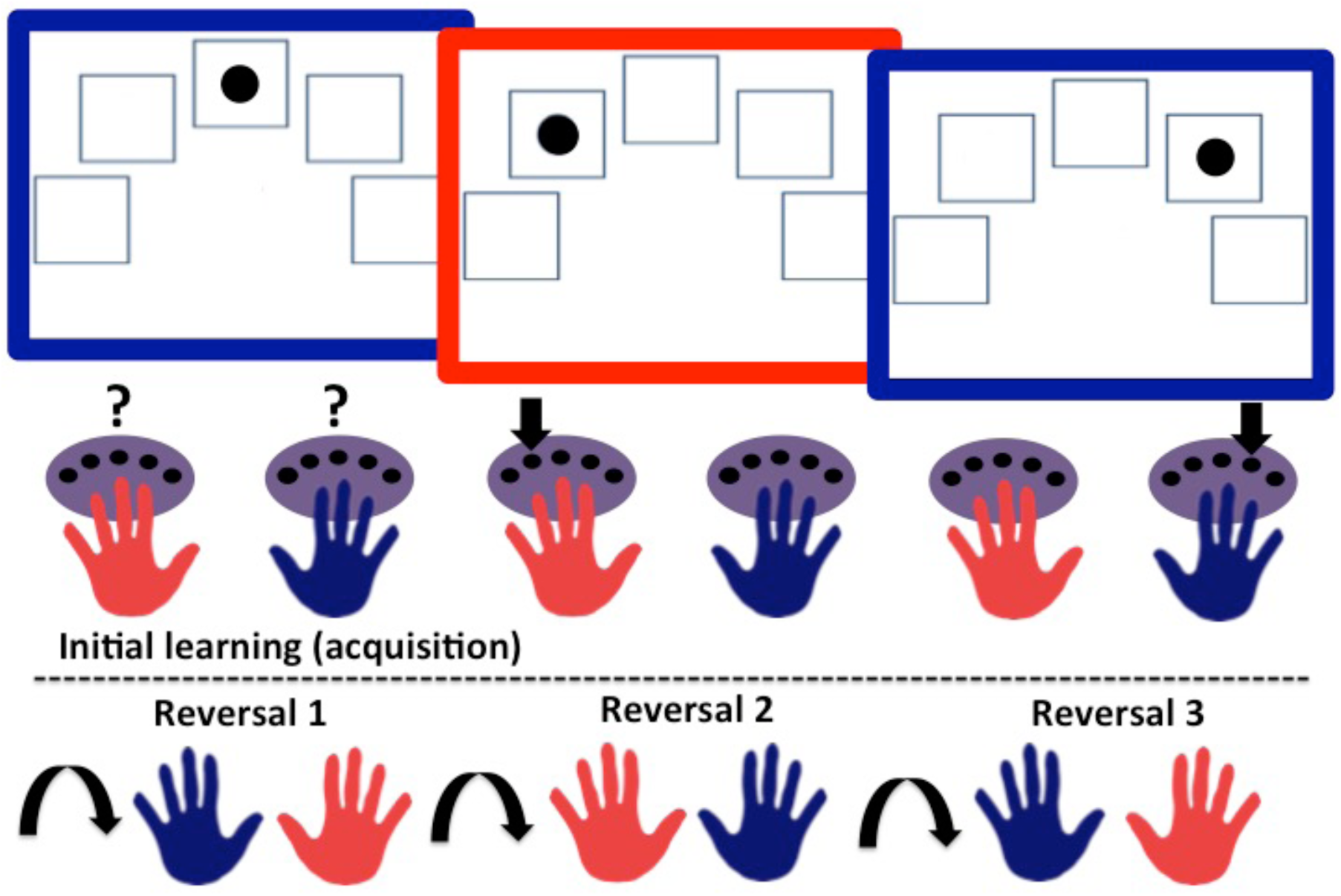
Experiment 1 task schematic. TOP: The three rectangles with coloured frames represent three example trials presented in the acquisition phase. Purple ovals symbolise the button boxes. Question marks signify the need to learn the correct hand-colour association by trial and error. Downward pointing arrows indicate the correct hand and button response for that trial. BOTTOM: Curved arrows signify the reversal of colour-hand contingencies, which occurred three times.

In the Pavlovian domain, meanwhile, it remains unknown how serotonin modulates reversal learning in any species. In previous human studies, researchers have focused on the influence of lowered serotonin signalling on initial Pavlovian threat conditioning (Hensman et al., 1991; Hindi Attar et al., 2012; Robinson et al., 2012). Here we tested whether instrumental reversal learning impairments additionally extend to the Pavlovian domain in humans following ATD (Figure 2). Whilst the primary hypotheses tested here pertained to the effects of serotonin on instrumental and Pavlovian reversal learning, we also expanded upon the few previous Pavlovian studies by examining threat conditioning to facial signs of aggression (innate threat cues; Ohman, 2009). Prior studies had paired initially neutral cues with aversive outcomes (e.g. heat or mild electric shock; Hensman et al., 1991; Hindi Attar et al., 2012; Robinson et al., 2012). Whilst similar neural mechanisms are engaged, there are distinctions between circuits that respond to learned threats (e.g. neutral cues), predators (e.g. snakes or spiders), and aggressive conspecifics (Gross & Canteras, 2012). Furthermore, in rodents, serotonin can be engaged differentially by innate versus learned threats (Isosaka et al., 2015) and by the intensity of threat (Seo et al., 2019).

**Figure 2.**
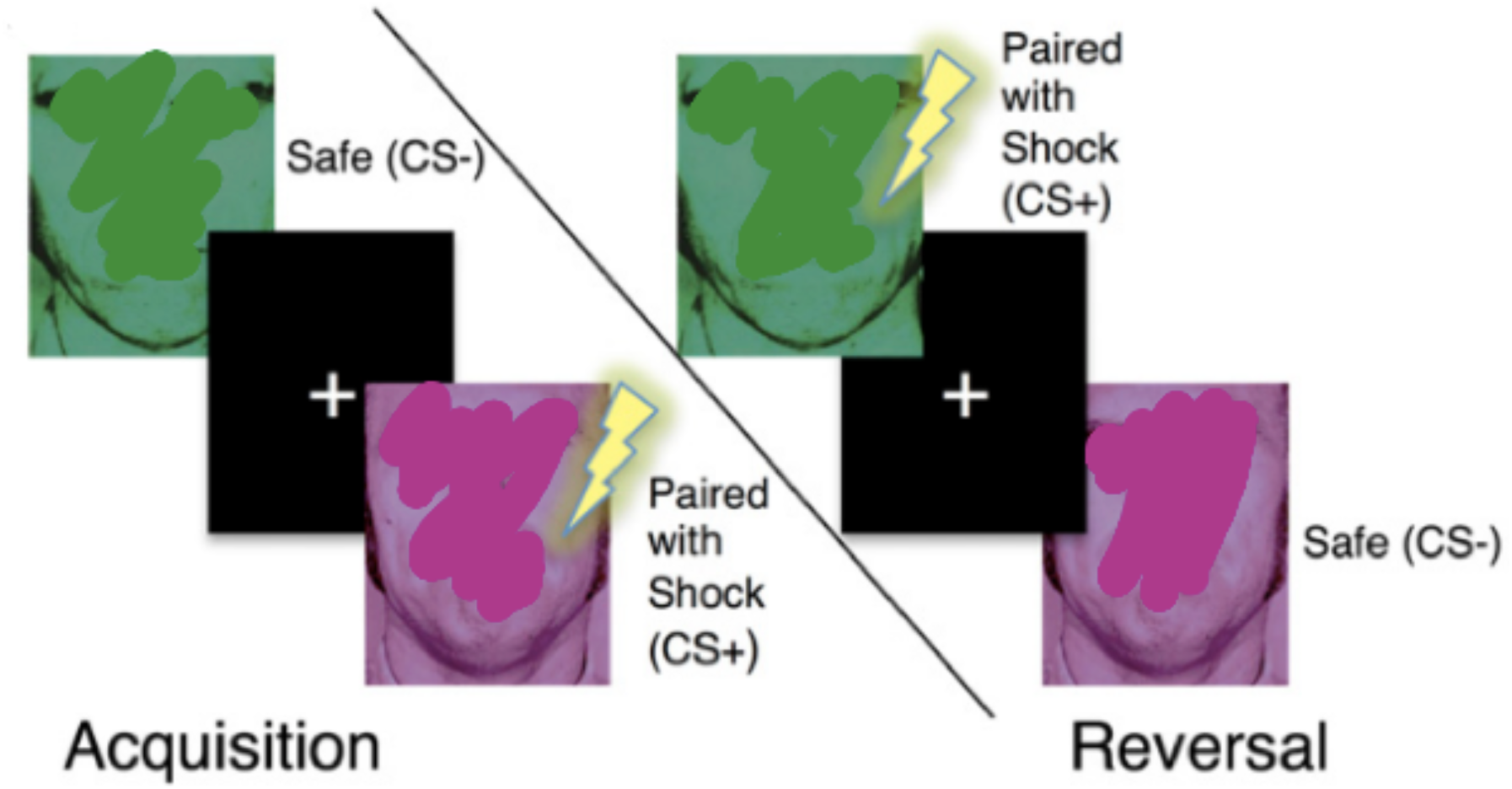
Experiment 2 task schematic, taken from Apergis-Schoute et al. (2017) with permission. Face stimuli were used with permission from Paul Ekman, PhD/Paul Ekman, LLC, and are altered for this pre-print Figure to comply with bioRxix’s policy of avoiding the inclusion of photographs of people.

The aim of this study, therefore, was to address the following questions. In Experiment 1: Does ATD induce perseveration in instrumental reversal learning? Do these effects emerge in a later reversal phase, and particularly when feedback is most salient? In Experiment 2: Does ATD impair Pavlovian reversal learning? And does ATD have a different effect on conditioning to threatening cues compared to neutral cues? In the instrumental domain, we hypothesised that ATD would lessen the impact of motivationally salient feedback to guide behaviour, resulting in a perseverative deficit. In the Pavlovian domain, we predicted conditioning processes would be impaired during both the initial acquisition phase as well as following the reversal of contingencies. Instrumental and Pavlovian reversal learning deficits following serotonin depletion would collectively point to a requirement of serotonin for integrating new information about reinforcement contingencies, which is fundamental to daily life and well-being.

## Results

### EXPERIMENT 1

#### Blood Results and Mood

Robust tryptophan depletion was achieved, as verified by plasma samples (t_(64)_ = −18.725, p = 1.1612 × 10^−27^). Plasma levels were unavailable for three participants: one due to a staff processing error, and two due to difficulty with venepuncture. Self-reported mood assessed using a visual analogue scale (VAS), available for 63 participants, was unaffected by ATD (t_(61)_ = -.898, p = .373).

#### Group-level instrumental learning

##### Omnibus analysis

Instrumental reversal learning was impaired following ATD, and the core deficits are displayed in Figure 3. First, omnibus repeated measures analysis of variance (ANOVA) was performed across all valence conditions and blocks. In the most salient condition participants had to make separate responses to obtain reward and avoid punishment (reward-punishment; see Methods). The other conditions incorporated either only neutral feedback (neutral-neutral), or neutral feedback with reward (reward-neutral) or punishment (punishment-neutral). The dependent measure for all analyses was trials to criterion (see Methods). The omnibus ANOVA, with serotonin status (placebo, depletion) as between-subjects factor, valence (reward-punishment, reward-neutral, punishment-neutral, neutral-neutral) and block (acquisition, reversal 1, reversal 2, reversal 3) as within-subjects factors, revealed a significant serotonin-by-valence-by-block interaction (F_(9,603)_ = 2.024, p = .035, η_p_^2^ = .029). There was no main effect of serotonin status (F_(1,67)_ = 1.869, p = .176, η_p_^2^ = .027).

**Figure 3.**
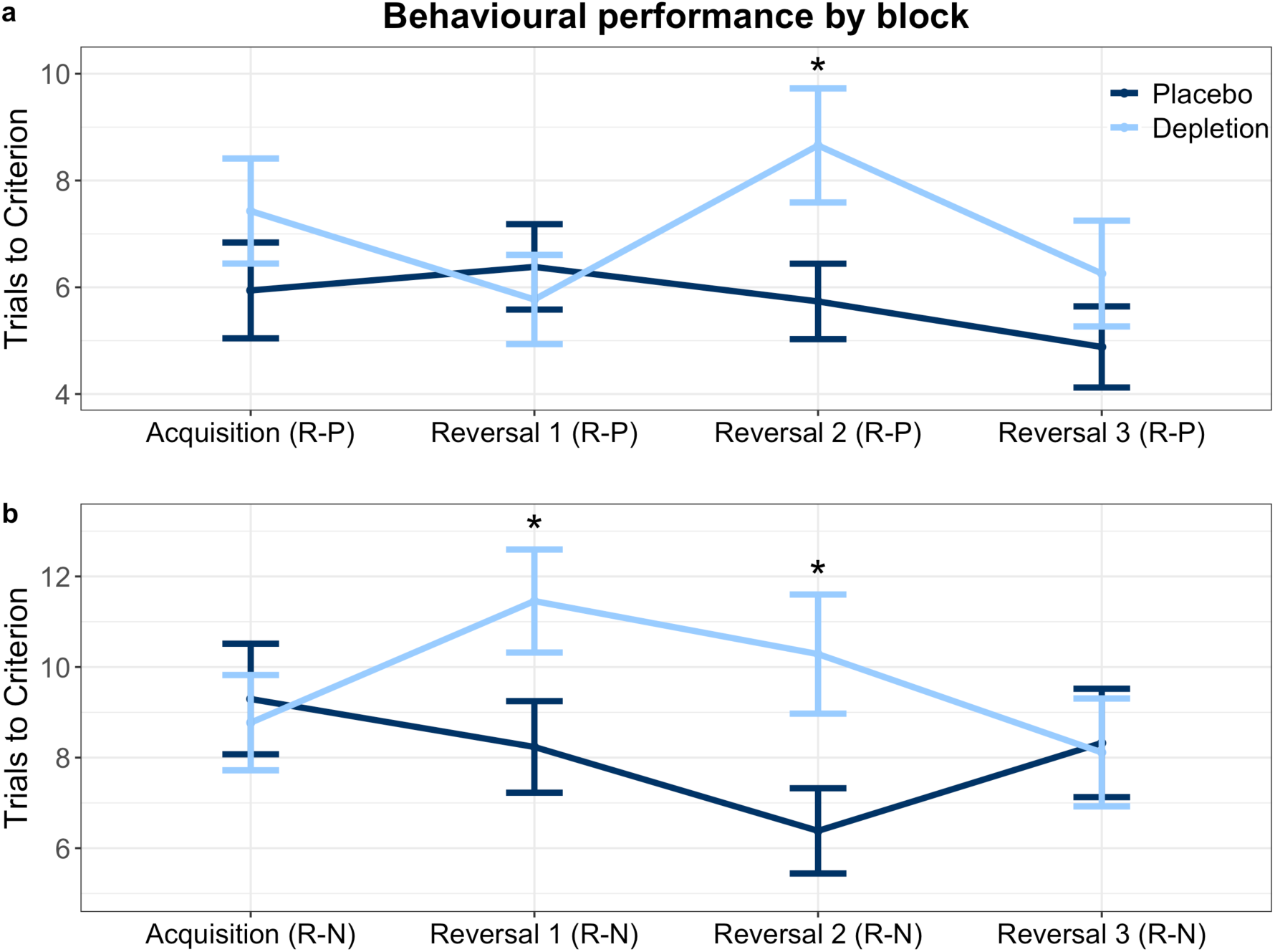
Instrumental reversal learning performance by block. More trials to criterion signifies worse performance. **(a)** Impairment in Reversal 2 of the reward-punishment condition = R-P; **(b)** Impairments in Reversals 1 and 2 of the reward neutral condition = R-N. Asterisks represent significance at p < .05. Error bars represent 1 standard error of the mean. There were no ATD-induced differences in the neutral-neutral condition or the punishment-neutral condition (not shown).

##### Acquisition learning

Next we verified that this effect was not driven by acquisition learning. Indeed, ATD had no effect on initial discrimination learning in the reward-punishment condition (t_(67)_ = 1.115, p = .269), reward-neutral (t_(67)_ = -.325, p = .746), punishment-neutral (t_(67)_ = -.688, p = .494) or neutral-neutral conditions (t_(64)_ = .891, p = .376), shown in Figure 3.

##### Reversal blocks

To assess the nature of the reversal learning deficit, the significant three-way interaction was followed up with t-tests in a sequence guided by two key a priori hypotheses. First, serotonin signalling is particularly engaged when responding to motivationally salient feedback (Faulkner and Deakin 2014), and therefore a reversal learning deficit should be most likely in the highest salience condition (reward-punishment). Second, serotonin depletion in the marmoset monkey OFC has been shown to induce the most pronounced instrumental reversal learning deficit in the second reversal block, without impacting the initial reversal (Clarke et al. 2004). The first follow-up test of reversal learning, therefore, assessed the second reversal of the most salient condition (reward-punishment) and indeed revealed a deficit: participants under ATD required more trials to criterion than on placebo (t_(59)_ = 2.281, p = .026). We then tested whether the effect in the second reversal was present in the other, less salient, conditions. There was a significant deficit under ATD in the reward-neutral condition (t_(61)_ = 2.413, p = .019), and not in the punishment-neutral (t_(67)_ = -.512, p = .61) or neutral-neutral (t_(67)_ = .572, p = .569) conditions. Next we tested whether a deficit was present in the first reversal. Individuals under depletion required more trials to criterion in the reward-neutral condition (t_(67)_ = 2.113, p = .038), but not in the reward-punishment (t_(67)_ = -.528, p = .599), punishment-neutral (t_(67)_ = 1.439, p = .155), or neutral-neutral (t_(64)_ = 1.051, p = .297) conditions. Finally, we assessed whether there was any deficit in the last reversal block. Performance was not impaired in the final reversal phase in the reward-punishment (t_(67)_ = 1.097, p = .277), reward-neutral (t_(67)_ = -.124, p = .902), punishment-neutral (t_(67)_ = 1.348, p = .182), or neutral-neutral (t_(67)_ = -.526, p = .601) conditions. The key deficits, from the reward-punishment and reward-neutral conditions identified in the second reversal block, additionally survived the Benjamini-Hochberg procedure (see Methods), for 12 comparisons (four valence conditions and three reversals), and were therefore the primary drivers of the serotonin-by-valence-by-block interaction.

#### Relationship between instrumental reversal deficits and extent of depletion

More pronounced depletion was significantly correlated with the key reversal deficits, shown in Figure 4. To further substantiate the deficits observed upon depletion, correlation analyses between behaviour and individual subject plasma samples were conducted. First, this was performed for behaviour in the second reversal block during both the reward-punishment and reward-neutral conditions, where significant deficits were found at the group level. Indeed, greater extent of depletion was significantly correlated with the magnitude of these key reversal impairments: more pronounced depletion was related to worse performance in both the reward-punishment condition (r_(66)_ = -.266, p = .031) and the reward-neutral condition (r_(66)_ = -.25, p = .043). These results are displayed in Figure 4a and 4b, respectively. The other observed behavioural impairment, from the first reversal in the reward-neutral condition, was also significantly correlated with the extent of depletion (r_(66)_ = -.311, p = .011).

**Figure 4.**
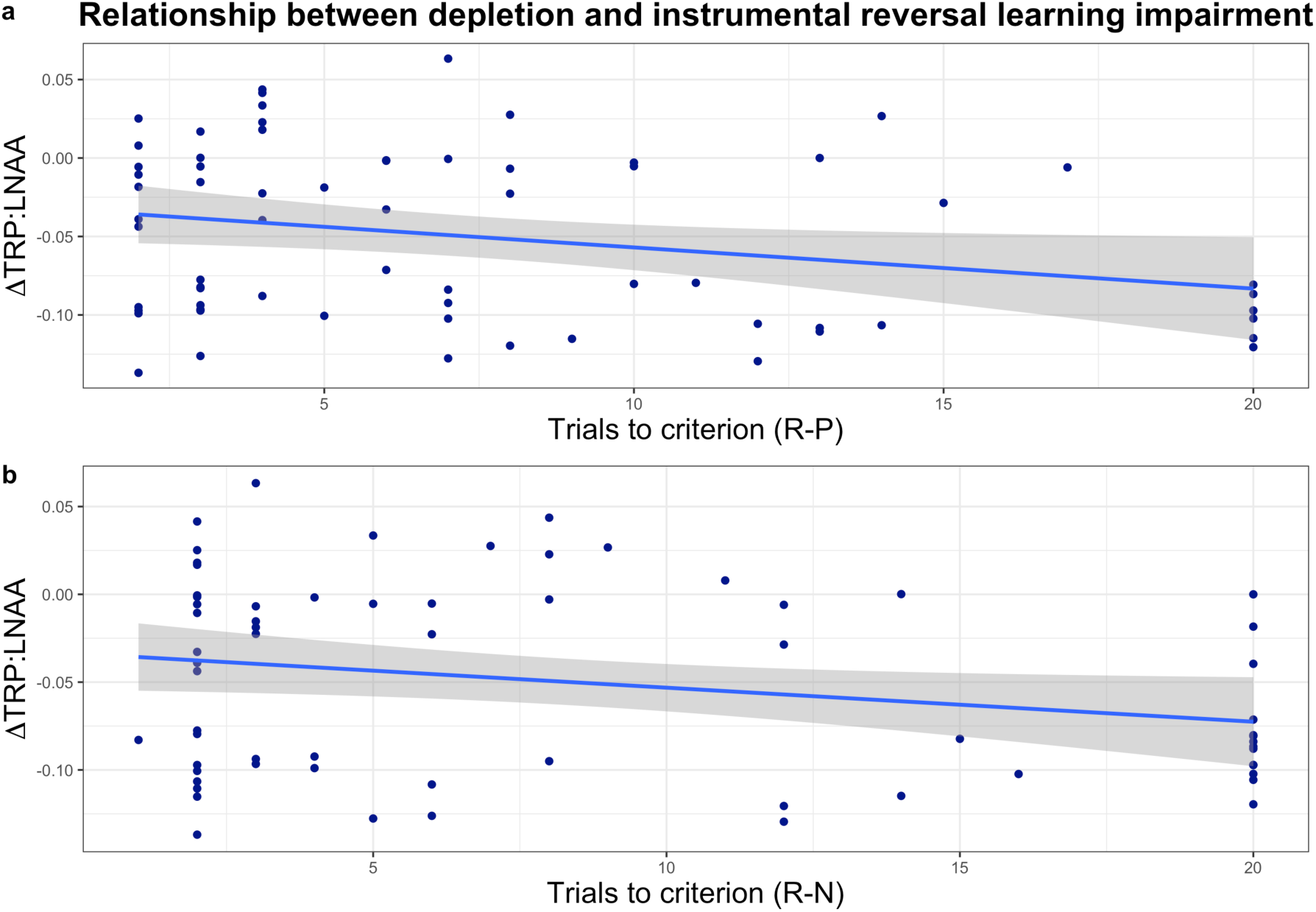
Relationship between extent of depletion and instrumental reversal learning performance. **(a)** Reward-punishment (R-P) condition. **(b)** Reward-neutral (R-N) condition. TRP:LNAA is the ratio of tryptophan to large neutral amino acids; y = 0 indicates no change. A greater decrease (post-depletion minus pre-depletion blood plasma results) in the ΔTRP:LNAA ratio indicates a more extensive depletion (more negative y-axis values). Reversal learning is indexed here as the number of trials to criterion in the second reversal block. Increasing x-axis values represent more trials to criterion and thus worse reversal performance. Shading indicates 1 standard error (SE).

### EXPERIMENT 2

#### Blood analysis and Mood

Robust depletion was also achieved in Experiment 2 (t_(17)_ = −4.907, p = 0.000132). Blood results from one participant were unavailable. Mood, assessed with the positive and negative affect schedule (PANAS; Watson et al. 1988) after depletion had taken effect, was unaffected: no difference between serotonin status for positive (t_(26)_ = 1.479, p = .151) or negative affect (t_(25)_ = −1.076, p = .292).

#### Acquisition of conditioning

Conditioning data are displayed in Figure 5a and 5b. Differential conditioning (CS+ versus CS-) was attained in both the placebo and ATD groups (paired t-tests: t_(11)_ = 6.866, p = .000027, for placebo; t_(15)_ = 7.181, p = .000003, for depletion). Conditioning was significantly stronger following depletion compared to the placebo group: we calculated a difference score of CS+ minus CS-for each group, and the magnitude of the CS+ relative to the CS-was significantly greater in the ATD group (t_(26)_ = −2.245, p = .034).

**Figure 5.**
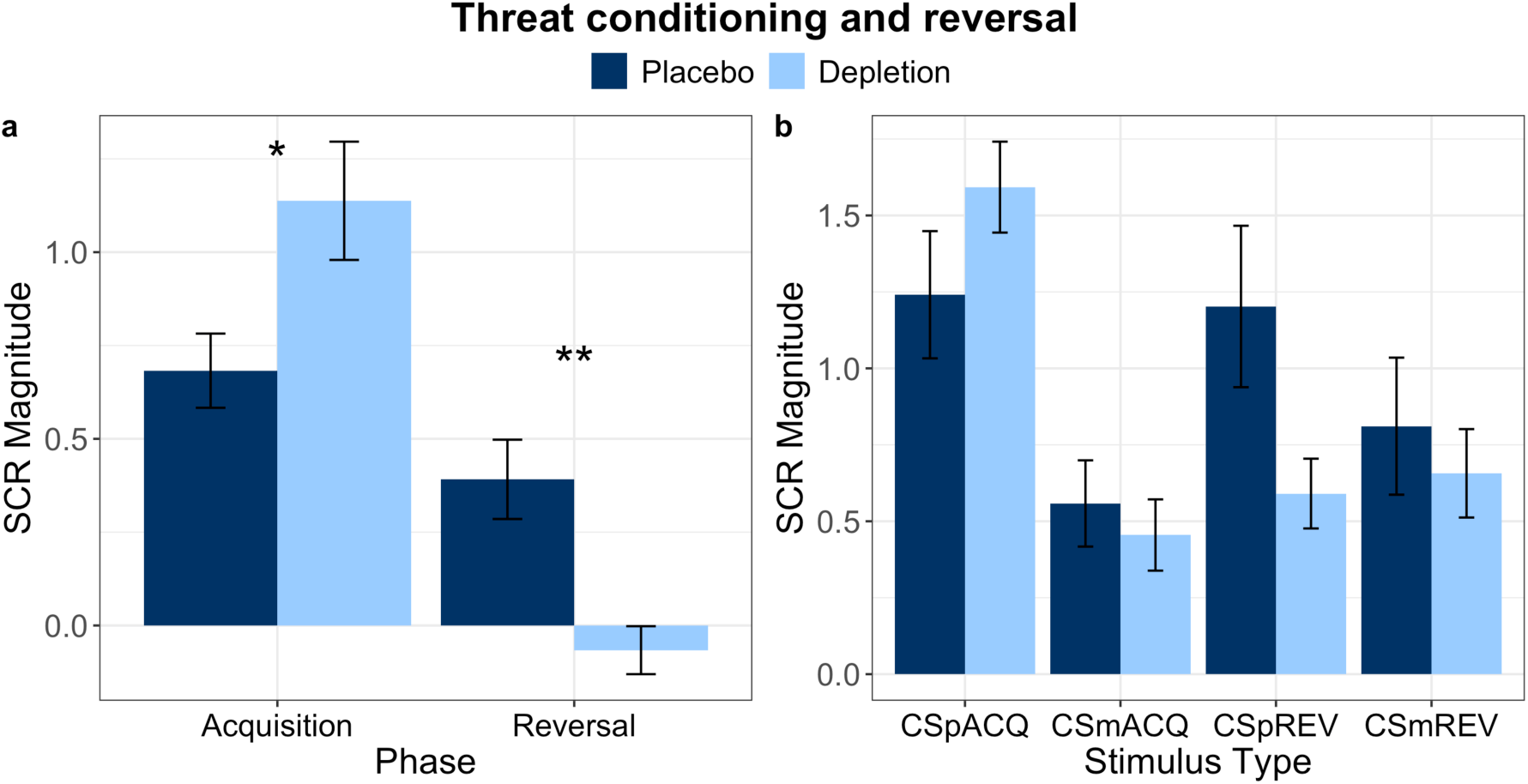
Pavlovian acquisition and reversal SCR data (Experiment 2), visualised in two different ways. Error bars represent 1 standard error (SE). **(a)** Difference scores of CS+ minus CS-, indicative of the extent of discrimination learning to the two stimuli. * indicates significance at p < .05; ** denotes significance at p < .01. **(b)** All stimuli displayed separately. CSpACQ = (initial) CS+ during acquisition; CSmACQ = (initial) CS-during acquisition; CSpREV = (new) CS+ during reversal; CSmREV = (new) CS-during reversal.

#### Reversal of conditioning

The reversal learning results are depicted in Figure 5a and 5b. During the reversal phase, the placebo group successfully conditioned to the new CS+ (t_(11)_ = 3.684, p = .004). The depletion group, however, did not show discrimination between the new CS+ and the new CS-(t_(15)_ = −1.031, p = .319), indicating a reversal learning impairment. Comparing the difference score during the reversal phase (new CS+ minus new CS-) between placebo and ATD also confirmed reversal learning was impaired (t_(26)_ = 3.880, p = .001).

### Changes in physiological responses to stimuli across phase

We additionally assessed how responses to the stimuli were affected by ATD in each phase, depicted in Figure 5b. Repeated measures analysis of variance (ANOVA), with serotonin status (placebo, depletion) as a between-subjects factor and phase (acquisition, reversal) and stimulus (CS+, CS-) as within-subjects factors revealed a significant three-way serotonin-by-phase-by-stimulus interaction (F_(1,26)_ = 17.604, p = .00028). Follow-up paired t-tests showed that responding to the initial CS+ extinguished upon reversal both within the placebo group (initial CS+ versus new CS-; t_(11)_ = 2.799, p = .017) and under ATD (t_(15)_ = 6.402, p = .000012). SCR to the initial CS-increased upon reversal in the placebo group (t_(11)_ = −4.172, p = .002) but critically, there was no difference in SCR to the initial CS-in acquisition compared to the new CS+ (old CS-) in reversal (t_(15)_ = −1.370, p = .191). The reversal impairment following ATD was driven by a failure to assign new aversive value, whereas safety learning upon reversal was intact.

#### Correlations between emotional measures and extent of depletion

Next we tested whether the extent of depletion, as assessed via plasma samples, was related to our measures of emotional learning. Extent of depletion was not correlated with the magnitude of the SCR difference score in the acquisition phase (r_(27)_ = .210, p = .294); however, there was a highly significant correlation between greater depletion and a more pronounced reversal learning deficit (r_(27)_ = -.536, p = .004), depicted in Figure 6. The Pavlovian reversal learning deficit was indexed by SCR to the CS+ minus the CS-in the reversal phase.

**Figure 6.**
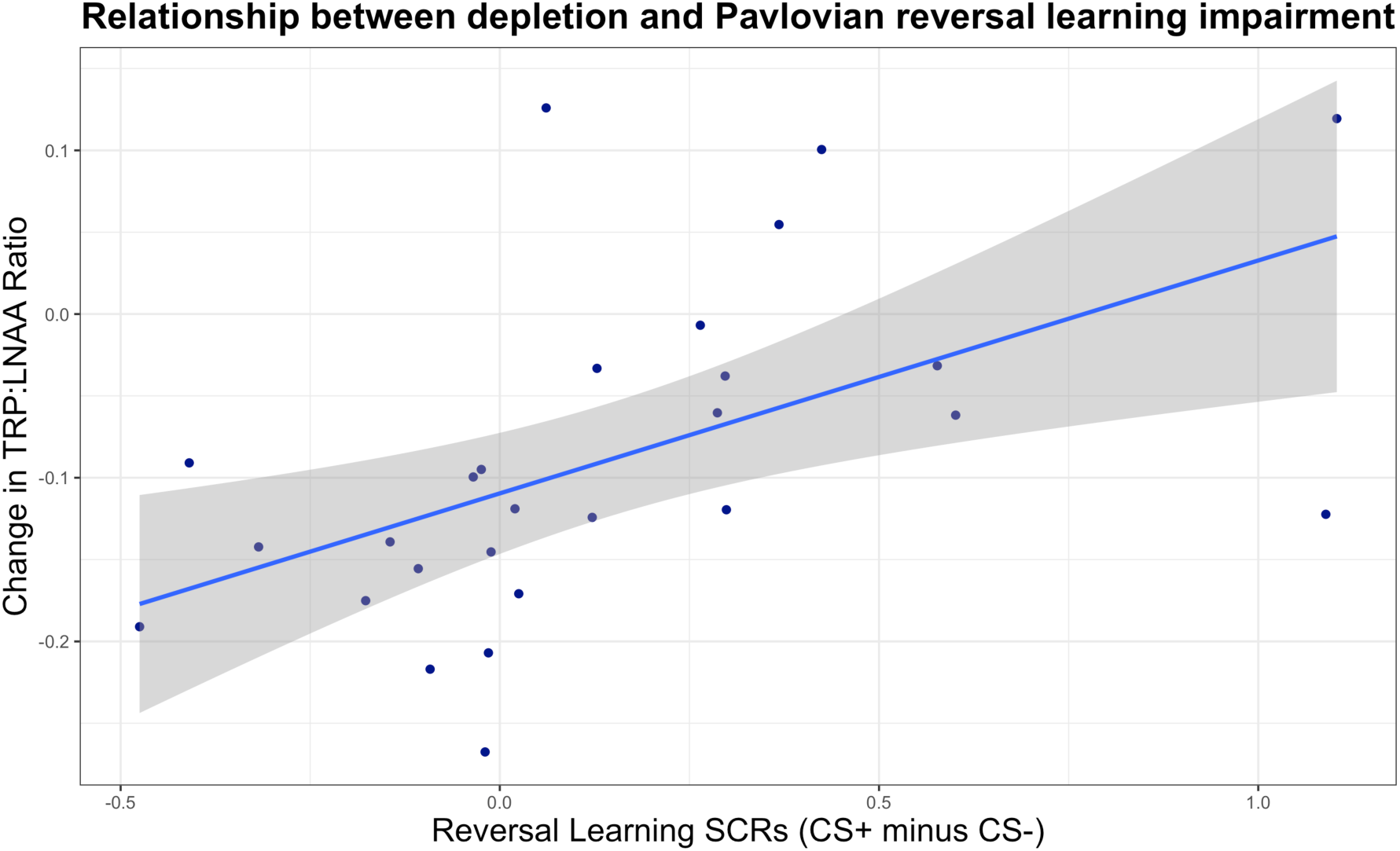
Experiment 2. Relationship between extent of depletion and degree of Pavlovian reversal learning impairment. TRP:LNAA is the ratio of tryptophan to large neutral amino acids; y = 0 indicates no change. A greater change (post-depletion blood minus pre-depletion results) in the TRP:LNAA ratio indicates a more extensive depletion (more negative y-axis values). Reversal learning is indexed here as the difference score between CS+ and CS-in the reversal phase. Increasing x-axis values represent better discrimination learning assessed by SCR between the CS+ and CS-in the reversal phase (i.e. better reversal learning). Shading indicates 1 standard error (SE).

## Discussion

We have provided convergent evidence from two independent experiments that serotonin depletion effected by acute dietary tryptophan depletion impairs human reversal learning in both the instrumental and Pavlovian domains (Experiments 1 and 2, respectively). The magnitude of the instrumental and Pavlovian reversal deficits, moreover, were both correlated with the extent of depletion assessed by plasma samples. Both the human instrumental and Pavlovian results are further strengthened by their consistency with studies of experimental animals following neurotoxic serotonin depletion (Bari et al. 2010; Clark et al. 2004, 2007; Lapiz-Bluhm et al. 2009). Remarkably, in rats, marmosets, and humans, the effect of serotonin depletion in the instrumental domain most consistently emerged upon the second reversal of contingencies (Bari et al. 2010; Clark et al. 2004, 2007). Pavlovian extinction, meanwhile, was intact following serotonin depletion in humans, which is also consistent with data from marmosets following OFC serotonin depletion: (instrumental) extinction was unimpaired (Walker et al. 2009). Initial Pavlovian conditioning here, to innately threatening cues, at the same time, was enhanced under serotonin depletion. This effect has not been seen for conditioning to innately neutral cues (Hensman et al. 1991; Hindi Attar et al. 2012). Mood was unaffected, in line with the ATD literature in healthy humans (Bell et al. 2005).

Perseverative deficits in human instrumental reversal learning following ATD have not been easily captured to date (Evers et al. 2005; Finger et al. 2007; Murphy et al. 2002), owing in part to ATD inducing a transient and relatively mild depletion in comparison (Worbe et al. 2014) with the profound depletion that is possible in experimental animals using 5,7-dihydroxytryptamine (Bari et al. 2010; Clarke et al. 2004, 2007; Walker et al. 2009). These ATD studies employed largely probabilistic feedback (Evers et al. 2005; Finger et al. 2007; Murphy et al. 2002), with a single reversal (Finger et al. 2007; Murphy et al. 2002), and non-salient feedback (Evers et al. 2005; Finger et al. 2007; Murphy et al. 2002). The innovative instrumental task used here was unique in that it incorporated highly salient feedback, multiple reversals on a deterministic schedule, and increased cognitive load. The deterministic schedule with multiple reversals, in particular, aligns with the design of marmoset studies that have provided quintessential evidence that OFC serotonin depletion induces perseveration (Clarke et al. 2004, 2007).

Whilst the instrumental deficits on both the most salient (reward and punishment) and reward-only, but not punishment-only condition, as reported here, may at first seem surprising given the well-established role of serotonin in aversive processing (Cools et al. 2008), this indeed aligns with the literature across species: the key marmoset studies on serotonin depletion and perseveration were conducted in the appetitive domain (Clarke et al. 2004; 2007; Walker et al. 2009), and Seymour et al. (2012) found human ATD affected the appetitive but not aversive domain in a 4-choice probabilistic task on which computational modelling revealed enhanced perseveration.

The Pavlovian reversal findings reported here resemble both the Pavlovian reversal learning impairment reported in OCD (Apergis-Schoute et al. 2017) and what has been demonstrated in healthy volunteers under stress (Raio et al. 2017). The reversal deficit in OCD, indexed by SCR on an identical paradigm, was explained by dysfunctional activity in the ventromedial prefrontal (vmPFC), which receives rich serotonergic innervation (Hornung 2010). Raio et al. (2017), meanwhile, used SCR and a similar design to that used here (but with neutral cues) and found that upon reversal, stress also attenuated the acquisition of threat responses to the newly threatening (previously safe) stimulus upon reversal, while leaving extinction learning to the previously threatening cue intact. This parallel is striking, and is consonant with data from rats: stress, and separately serotonin depletion, produced comparable deficits in (instrumental) reversal learning (Lapiz-Bluhm et al. 2009). Serotonin release in rats during behavioural testing, moreover, was reduced by stress, and an SSRI given acutely ameliorated the detrimental effect of stress on reversal learning (Lapiz-Bluhm et al. 2009).

The deleterious effects of serotonin depletion and stress on reversal learning can be interpreted as a selective impairment in integrating new information about a change in reinforcement contingencies, needed to update the representation of aversive value appropriately (Raio et al. 2017). These authors invoked the phenomenon of stress-induced dopamine release (Pruessner et al. 2004), which may dampen negative prediction errors evoked by contingency reversal (Cools et al. 2001). Our reversal data are consistent with a similar interpretation: ATD may have constrained the negative prediction errors triggered by reversal, which are purported to be associated with serotonin signalling (Cools et al. 2011).

That we found an enhancement of initial Pavlovian conditioning, when employing socially threatening conditioned stimuli (aggressive faces), contrasts with two previous threat conditioning studies that used neutral stimuli: Hensman et al. (1991) showed that the serotonin 2A/2C (5-HT2A, 5-HT2C) receptor antagonist ritanserin attenuated threat conditioning, whilst Hindi Attar et al. (2012) showed the same pattern of results following ATD. The attenuated SCR upon reversal as reported here, however, converges with these previous studies that investigated conditioning alone (Hensman et al. 1991; Hindi Attar et al. 2012). The discrepancy across studies at the initial conditioning phase (Hensman et al. 1991; Hindi Attar et al. 2012) is likely due to the engagement of distinct, yet similar, mechanisms when learning to predict aversion from a neutral cue, compared to responding to innate threats (Gross & Canteras 2012).

Consideration of rodent studies on the influence of serotonin on specific amygdala sub-nuclei may inform our human conditioning findings. The central nucleus of the amygdala (CeA) is the major source of output and its downstream projections ultimately produce defence responses such as perspiration in humans and freezing in rodents (LeDoux 2000). Critically, cells expressing 5-HT2A receptors in the CeA are differentially engaged by innate versus learned threats (Isosaka et al. 2015). Inhibition of these 5-HT2A-expressing cells upregulates innate threat responses in mice and downregulates learned threat responses (Isosaka et al. 2015). This is remarkably congruent with our observation that reducing serotonin function potentiates conditioning to innate threats, on the one hand, and findings from previous studies that reduction of serotonin signalling attenuates threat conditioning to learned (neutral) cues. These divergent results may inform therapeutic, and possibly adverse, effects of serotonin modulating drugs. Isosaka et al. (2015) indeed showed that risperidone exacerbated responses to innate threats and alleviated threat responses to previously neutral cues: this latter result is consistent with Hensman et al. (1991). Furthermore, humans with selective damage to the basolateral amygdala (BLA), with the CeA preserved, showed hypervigilant responses to fearful faces (innate threat), which was interpreted as the removal of an inhibitory influence of the BLA over the CeA (Terburg et al. 2012). Indeed, the BLA receives particularly rich serotonergic innervation (Bauer 2015), which, in conjunction with the role of serotonin in the CeA for innate threats may be important for understanding our results.

As in other human ATD studies we did not measure serotonin directly, and instead used a widely accepted proxy measurement (Bell et al., 2005). Whilst some have criticised ATD as a technique for studying serotonin in particular (van Donkelaar et al., 2011), the method has been robustly defended (Crockett et al., 2012a). Critically, the present findings align with deficits following profound neurotoxic serotonin depletion in both rats (Bari et al. 2010) and marmoset monkeys (Clarke et al. 2004, 2007; Walker et al. 2009). Our results build upon other studies, for instance on “waiting impulsivity”, that show parallel behavioural effects between neurotoxic depletions in experimental animals (Winstanley et al., 2004) and ATD in healthy humans (Worbe et al., 2014), thus further bolstering the validity of ATD for studying serotonin. Whilst we don’t know the neural locus of this impaired neuromodulation, work in the instrumental domain from experimental animals (Clarke et al. 2004, 2007; Walker et al. 2009) and individuals with OCD (Chamberlain et al. 2008; Remijnse et al. 2006) enable us to highlight the OFC. The Pavlovian reversal data from OCD, meanwhile, point to the vmPFC (Apergis-Schoute et al. 2017).

We provide evidence of human reversal learning impairments following serotonin depletion, in both the instrumental and Pavlovian domains, across two independent experiments. Deficits in both domains were underscored by significant correlations showing that a greater extent of depletion, as assessed by plasma samples, was associated with more pronounced reversal impairments. Strikingly, the results align with data from neurotoxic serotonin depletion in experimental animals (Bari et al. 2010; Clarke et al. 2004, 2007; Walker et al. 2009), stress induction in humans (Raio et al. 2017) and rats (Lapiz-Bluhm et al. 2009), and individuals with OCD (Apergis-Schoute et al. 2017).

That serotonin depletion impaired these fundamental learning processes pervasive in daily life highlights a failure mode that could lead to significant distress and impairment. The reversal deficits presented, furthermore, indicate how serotonergic dysfunction could impede the ability to engage in cognitive behavioural therapies (CBT). The present results therefore advance knowledge on the neurochemical basis of emotional and behavioural flexibility, which has implications for the understanding and treatment of numerous clinical conditions including OCD.

## Methods

### EXPERIMENT 1

#### Participants

Sixty-nine healthy participants (36 males, mean age 24.28) completed the deterministic reversal learning task and were included in the final analysis. One male participant in the depletion group was excluded because he admitted to responding randomly later in the task. Participants were screened to be medically healthy and free from any psychiatric conditions, determined by the Mini International Neuropsychiatric Interview (MINI; Sheehan, et al., 1998). Individuals who reported having a first-degree relative (parent or sibling) with a psychiatric disorder were excluded upon screening as well. Exclusion criteria also encompassed neurological disorders; pregnancy; past use of neurological, psychiatric, or endocrine medication (including St John’s wort); or current use of any regular medication besides contraceptive pills. The cut-offs for drug use were smoking more than five cigarettes per day, regular consumption of more than 38 UK units (380 ml) of alcohol per week, cannabis use more than once per month, and the lifetime use of recreational drugs besides cannabis more than five times. Other medical exclusion criteria: cardiac or circulation issues, respiratory problems including asthma; gastrointestinal, renal, or thyroid conditions; bleeding disorders, diabetes and head injury. Volunteers provided informed consent before the study and were paid for their participation. Groups did not differ in age, years of education, trait impulsivity, or in baseline depressive and obsessive-compulsive symptoms, shown in Supplementary Table 1.

#### General Procedure

The protocol was approved by the Cambridge Central NHS Research Ethics Committee (Reference # 16/EE/0101). The study took place at the National Institute for Health Research / Wellcome Trust Clinical Research Facility at Addenbrooke’s Hospital in Cambridge, England. Participants arrived in the morning having fasted for at least 9 hours prior, gave a blood sample, and ingested either the placebo or ATD drink. To assess mood and other feelings including alertness, we used a 16-item visual analogue scale (VAS) at the beginning, middle, and end of the day-long testing session. In the afternoon participants completed the deterministic reversal learning task, along with several other tasks that will be reported elsewhere.

#### Acute Tryptophan Depletion

Tryptophan, the precursor required for serotonin synthesis, is an essential amino acid: it cannot be produced by the body and therefore must be obtained from the diet. Acute tryptophan depletion (ATD) is therefore an ingenious dietary technique for the study of serotonin, which rapidly decreases serotonin function (Bel & Artigas, 1996; Biggio et al., 1974; Crockett et al., 2012a; Nishizawa et al. 1997). Healthy volunteers were assigned to receive ATD or placebo, in a randomised, double-blind, between-groups design. The ATD group consumed a drink containing the essential amino acids less tryptophan, whereas the placebo drink was identical other than it included tryptophan. Blood samples were taken to verify depletion.

#### Instrumental reversal learning task

The task used in Experiment 1 is depicted in Figure 1. As an incentive, participants were told that depending on how well they performed the task, they could win a bonus on their compensation for taking part in the study. In reality, everyone received a small bonus. The instrumental reversal paradigm was designed to increase cognitive load and thus task demands. It had three reversals and a deterministic schedule. Responses were entered via one of two “button boxes” with either the left or right hand, see Figure 1. On each trial, the computer screen was framed by a specific colour and displayed five boxes corresponding to five buttons on each button box, one button per finger. The colour indicated the correct hand to respond with, and a black dot inside one of the five boxes on the screen indicated which finger to respond with, depicted in Figure 1. Participants were told they needed to learn the colour-hand association by trial and error and that the association would change multiple times within a run. A run consisted of four blocks of 20 trials each: an acquisition block where the initial contingency was established followed by three reversal blocks. The reinforcement schedule was deterministic: the correct option led to positive feedback on 100% of trials, whilst the incorrect response led to negative feedback on 100% of trials. Trial order was randomised. There were four runs in random order, and each contained a unique pair of colours framing the screen which was counterbalanced. All runs contained the same visual feedback cartoon stimuli (shown in Supplementary Figure 1): a smiling face with “two thumbs up” for correct responses, a face showing disappointment and a “thumbs down” when incorrect, and an analogue alarm clock with a frown if a response was not entered within the allotted time. The salience and valence of feedback across runs was varied using the presence or absence of prominent auditory stimuli. The primary run of interest had the most salient auditory feedback: responding correctly to one colour resulted in reward in the form of a prominent “cha-ching” (slot machine) sound, whilst correct responses to the other colour prevented (avoided) the occurrence of an aversive buzzer noise (reward-punishment run). There was also a reward-neutral run where a correct response to one colour frame resulted in the reward auditory feedback whereas responding correctly or incorrectly to the other colour resulted only in visual (neutral) but no auditory feedback. In the punishment-neutral run incorrect responses to one colour frame were punished with the buzzer noise whereas correct or incorrect responses to the other colour resulted only in visual feedback (neutral). Finally, the task contained a neutral-neutral condition where no auditory feedback was provided and only visual feedback via cartoons was presented.

The experiment began with three training phases, each of which required making correct responses on at least 80% of trials to advance to the next stage otherwise the phase would be repeated. The first was self-paced and served to familiarise participants with responding using the button boxes. In the first training phase only, “LEFT” or “RIGHT” was displayed on each trial to instruct the correct hand to use. There was a time limit in the second (short) and third (longer) training phases and participants were told to respond as quickly and accurately as possible. The time window to make responses during the actual experiment was automatically calibrated to each person based on their reaction times during the final practice phase. The task was programmed in E-Prime 2.0 Professional. The primary dependent measure was trials to criterion, which has been used in serotonin depletion studies in marmoset monkeys (Clarke et al., 2004, 2007), which we aimed to translate into humans here. The criterion was defined as making four consecutive correct responses.

### EXPERIMENT 2

#### Participants

Thirty healthy volunteers (17 females) completed the Pavlovian threat reversal task. Of these, two (1 female) were “non-responders” and were thus excluded for an undetectable skin conductance response as is standard practice (e.g. Schiller et al. 2013). All participants provided written informed consent and were financially compensated. The study was approved by the Cambridgeshire 2 Research Ethics Committee (Reference # 09/H0308/51). Participants were eligible if they did not have a personal or family history of major depressive disorder, bipolar disorder, or any other psychiatric illness. Other exclusion criteria included medication use, a history of neurological, cardiac, gastrointestinal, hepatic, pulmonary, or renal disorders. Groups did not differ in age, years of education, trait impulsivity, or in baseline depressive symptoms, shown in Supplementary Table 2.

#### General procedure

Participants were assigned to receive either placebo or the tryptophan-depleting drink in a randomized, double-blind design (16 received depletion). Blood samples were collected at baseline and before the task to verify tryptophan depletion. Participants completed questionnaires including one assessing self-reported mood state. Data were collected inside of a functional magnetic resonance imaging (fMRI) scanner, but the fMRI data are not reported here. Participants returned for a second session, where they received the other drink condition and also completed different computerised tasks, results of which have been published elsewhere (Crockett et al., 2013; Passamonti et al., 2012). It is important that participants are naïve to conditioning paradigms, and therefore the data reported here were acquired in the first of two testing sessions spaced at least one week apart.

#### Conditioning procedure

The task is depicted in Figure 2 and had two phases: acquisition and reversal. Two faces (face A and B) were presented in each phase, for four seconds each with an inter-trial interval of 12 seconds (Schiller et al. 2008). The face images were selected from the Ekman series (Ekman & Friesen, 1976). Participants chose a shock level that they felt was uncomfortable but not painful. In the acquisition phase, face A was presented 16 times without a shock (conditioned stimulus plus; CS+) and coterminated with a 200 millisecond shock (unconditioned stimulus; US) on an additional eight trials (CS+US), while face B was presented 16 times and never paired with shock (CS-). In the reversal phase the faces were presented again only the contingencies swapped: face A was presented for 16 trials and was no longer paired with a shock (new CS-), while face B was newly paired with a shock on 8 trials amidst an additional 16 unreinforced trials (new CS+). Trials were pseudorandomized and designation of face A and B was counterbalanced. Reversal was unsignaled and immediately followed acquisition without a break. The dependent measure was the skin conductance response (SCR), which assesses perspiration and is an assay of autonomic nervous system arousal. The primary focus was the SCR to unreinforced trials, to avoid contamination by the shock itself. SCRs were defined as the base-to-peak difference during a seven second interval beginning 0.5 seconds after stimulus onset. SCRs were normalised for each individual participant by dividing values from each trial by the peak amplitude. More information about this specific paradigm can be found in Apergis-Schoute et al. (2017).

## Acknowledgements

This research was funded by a Wellcome Trust Senior Investigator Award (104631/Z/14/Z) to T.W.R. B.J.S. receives funding from the National Institute for Health Research (NIHR) Cambridge Biomedical Research Centre (Mental Health Theme); the views expressed are those of the authors and not necessarily those of the NIHR or the Department of Health and Social Care. R.N.C.’s research is supported by the UK Medical Research Council (MC_PC_17213). J.W.K. and M.J.C were supported by Gates Cambridge Scholarships. We would like to thank the staff at the NIHR/Wellcome Trust Clinical Research Facility at Addenbrooke’s Hospital and at the Wolfson Brain Imaging Centre where the experiments were conducted, and Rachel Kyd of the Cambridge University Hospital Research & Development Office for assistance with study approval.

## Supplemental Information

### Methods

Plasma samples were analysed using high performance liquid chromatography (HPLC) as in Crockett et al. (2013) and Passamonti et al. (2012). Depletion was indexed using the ratio of tryptophan to large neutral amino acids, thought to be most reflective of brain serotonin (Bell et al. 2005). The Greenhouse-Geisser correction was used where applicable, in designs with within-subjects factors, to correct for violation of the sphericity assumption as determined by Mauchly’s test. Degrees of freedom were adjusted for t-tests when Levene’s Test for equality of variances was violated. The critical value for the Benjamini-Hochberg procedure for false discovery rate was set at q = .15 per Skandali et al. (2018).

**Supplemental Table 1.** Demographics and questionnaire measures for Experiment 1. BDI-II = Beck Depression Inventory, version II (Beck et al. 1996); OCI-R = Obsessive Compulsive Inventory (Foa et al. 2002); BIS = Barratt Impulsiveness Scale (Patton et al. 1995). SD = standard deviation.

| **Group** | **Placebo Mean (SD)** | **Depletion Mean (SD)** | **t(df)** | **p-value** |
| --- | --- | --- | --- | --- |
| Age | 24.24 (5.684) | 24.31 (4.788) | .063 (67) | .95 |
| Years of education | 16.97 (2.181) | 17.34 (2.376) | .678 (67) | .5 |
| BDI-II | 4.76 (4.335) | 3.80 (3.701) | -.995 (67) | .323 |
| OCI-R | 8.21 (7.543) | 7.09 (7.172) | -.632 (67) | .529 |
| BIS | 70.41 (4.150) | 71.09 (3.673) | .715 (67) | .477 |

**Supplemental Table 2.** Demographics and questionnaire measures for Experiment 2. BDI-II = Beck Depression Inventory, version II (Beck et al. 1996); BIS = Barratt Impulsiveness Scale (Patton et al. 1995). SD = standard deviation. Age and years of education were unavailable for one participant in the placebo condition. In the depletion condition, age was unavailable for one participant; years of education were unavailable for two participants.

| **Group** | **Placebo Mean (SD)** | **Depletion Mean (SD)** | **t(df)** | **p-value** |
| --- | --- | --- | --- | --- |
| Age | 23.58 (8.262) | 25.21 (2.860) | -.694 (24) | .495 |
| Years of education | 18.33 (1.923) | 18.46 (2.727) | -.135 (23) | .894 |
| BDI-II | 3.42 (3.423) | 4.06 (4.739) | -.400 (26) | .693 |
| BIS | 55.33 (6.827) | 59.81 (8.272) | -1.524 (26) | .139 |

**Supplementary Figure 1.**
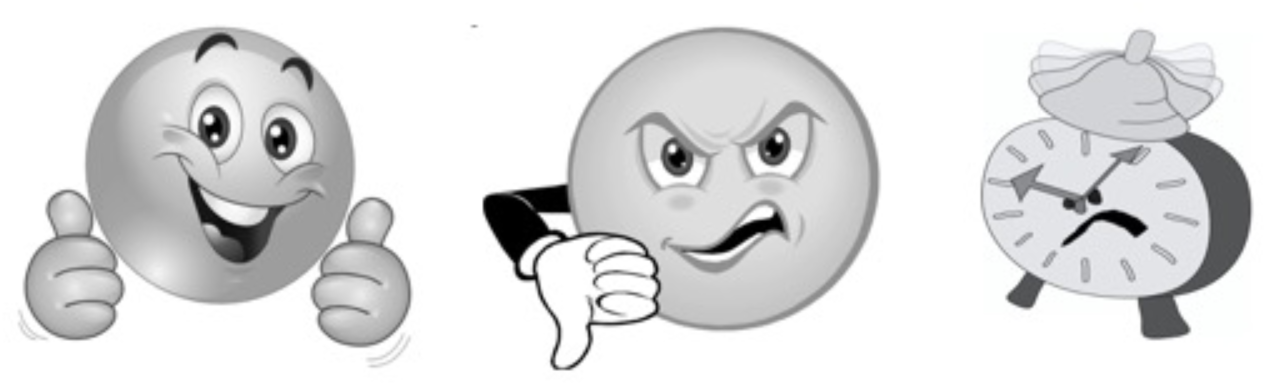
Feedback icons for the task. Left to right: correct, incorrect, too late.

## References

American Psychiatric Association (2013) Diagnostic and Statistical Manual of Mental Disorders, 5th Edition. American Psychiatric Association, Washington, DC

Apergis-Schoute AM, Gillan CM, Fineberg NA, et al (2017) Neural basis of impaired safety signaling in Obsessive Compulsive Disorder. Proc Natl Acad Sci U S A 114:3216–3221. doi: 10.1073/pnas.1609194114

Baldwin DS, Anderson IM, Nutt DJ, et al (2014) Evidence-based pharmacological treatment of anxiety disorders, post-traumatic stress disorder and obsessive-compulsive disorder: A revision of the 2005 guidelines from the British Association for Psychopharmacology. J Psychopharmacol 28:403–439. doi: 10.1177/0269881114525674

Bari A, Theobald DE, Caprioli D, et al (2010) Serotonin Modulates Sensitivity to Reward and Negative Feedback in a Probabilistic Reversal Learning Task in Rats. Neuropsychopharmacology 35:1290–1301. doi: 10.1038/npp.2009.233

Barlow RL, Alsiö J, Jupp B, et al (2015) Markers of Serotonergic Function in the Orbitofrontal Cortex and Dorsal Raphé Nucleus Predict Individual Variation in Spatial-Discrimination Serial Reversal Learning. Neuropsychopharmacology 40:1619–1630. doi: 10.1038/npp.2014.335

Bauer EP (2015) Serotonin in fear conditioning processes. Behav Brain Res 277:68–77. doi: 10.1016/j.bbr.2014.07.028

Beck AT, Steer RA, Ball R, Ranieri WF (1996) Comparison of Beck Depression Inventories-IA and-II in Psychiatric Outpatients. J Pers Assess 67:588–597. doi: 10.1207/s15327752jpa6703_13

Bel N, Artigas F (1996) Reduction of serotonergic function in rat brain by tryptophan depletion: Effects in control and fluvoxamine-treated rats. J Neurochem 67:669–676. doi: 10.1046/j.1471-4159.1996.67020669.x

Bell C, Hood S, Nutt D (2005) Acute tryptophan depletion. Part II: clincal effects and implications. Aust N Z J Psychiatry 39:565–574

Biggio G, Fadda F, Fanni P, et al (1974) Rapid depletion of serum tryptophan, brain tryptophan, serotonin and 5-hydroxyindoleacetic acid by a tryptophan-free diet. Life Sci 14:1321–1329. doi: 10.1016/0024-3205(74)90440-8

Chamberlain SR, Menzies L, Hampshire A, et al (2008) Orbitofrontal Dysfunction in Patients with Obsessive-Compulsive Disorder and Their Unaffected Relatives. Science (80-) 321:421–422. doi: 10.1126/science.1154433

Clarke HF, Dalley JW, Crofts HS, et al (2004) Cognitive Inflexibility After Prefrontal Serotonin Depletion. Science (80-) 304:878–880. doi: https://doi.org/10.1126/science.1094987

Clarke H., Walker SC, Dalley JW, et al (2007) Cognitive Inflexibility after Prefrontal Serotonin Depletion Is Behaviorally and Neurochemically Specific. Cereb Cortex 17:18–27. doi: 10.1093/cercor/bhj120

Cohen JY, Amoroso MW, Uchida N (2015) Serotonergic neurons signal reward and punishment on multiple timescales. Elife 2015:1–25. doi: 10.7554/eLife.06346

Cools R, Barker RA, Sahakian BJ, Robbins TW (2001) Enhanced or Impaired Cognitive Function in Parkinson’s Disease as a Function of Dopaminergic Medication and Task Demands. Cereb Cortex 11:1136–1143. doi: 10.1093/cercor/11.12.1136

Cools R, Roberts AC, Robbins TW (2008) Serotoninergic regulation of emotional and behavioural control processes. Trends Cogn Sci 12:31–40. doi: 10.1016/j.tics.2007.10.011

Crockett MJ, Clark L, Roiser JP, et al (2012a) Converging evidence for central 5-HT effects in acute tryptophan depletion. Mol Psychiatry 17:121–123. doi: 10.1038/mp.2011.106

Crockett MJ, Apergis-Schoute A, Herrmann B, et al (2013) Serotonin modulates striatal responses to fairness and retaliation in humans. J Neurosci 33:3505–13. doi: 10.1523/JNEUROSCI.2761-12.2013

Crockett MJ, Clark L, Apergis-Schoute AM, et al (2012b) Serotonin Modulates the Effects of Pavlovian Aversive Predictions on Response Vigor. Neuropsychopharmacology 37:2244–2252. doi: 10.1038/npp.2012.75

Cunningham KA, Howell LL, Anastasio NC (2020) Serotonin neurobiology in cocaine use disorder. In: Muller C, Cunningham KA (eds) Handbook of the Behavioral Neurobiology of Serotonin, 2nd Ed. Academic Press, pp 745–802

Dayan P, Huys QJM (2009) Serotonin in Affective Control. Annu Rev Neurosci 32:95–126. doi: 10.1146/annurev.neuro.051508.135607

Deakin JFW (2013) The origins of “5-HT and mechanisms of defence” by Deakin and Graeff: a personal perspective. J Psychopharmacol 27:1084–9. doi: 10.1177/0269881113503508

Dersken M, Feenstra M, Willuhn I, Denys D (2020) The serotonergic system in obsessive-compulsive disorder. In: Muller C, Cunningham KA (eds) Handbook of the Behavioral Neurobiology of Serotonin, 2nd Ed. Academic Press, pp 865–892

Ekman P, Friesen WV (1976) Pictures of Facial Affect (Consulting Psychologists Press, Palo Alto, CA)

Evers EA, Cools R, Clark L, et al (2005) Serotonergic modulation of prefrontal cortex during negative feedback in probabilistic reversal learning. Neuropsychopharmacology 30:1138–1147. doi: 10.1038/sj.npp.1300663

Faulkner P, Deakin JFW (2014) The role of serotonin in reward, punishment and behavioural inhibition in humans: Insights from studies with acute tryptophan depletion. Neurosci Biobehav Rev 46P3:365–378. doi: 10.1016/j.neubiorev.2014.07.024

Fineberg NA, Reghunandanan S, Brown A, Pampaloni I (2013) Pharmacotherapy of obsessive-compulsive disorder: Evidence-based treatment and beyond. Aust. N. Z. J. Psychiatry 47:121–141

Finger EC, Marsh A a, Buzas B, et al (2007) The impact of tryptophan depletion and 5-HTTLPR genotype on passive avoidance and response reversal instrumental learning tasks. Neuropsychopharmacology 32:206–215. doi: 10.1038/sj.npp.1301182

Foa EB, Huppert JD, Leiberg S, et al (2002) The Obsessive-Compulsive inventory: Development and Validation of a Short Version. Psychol Assess 14:485–496. doi: 10.1037/1040-3590.14.4.485

Gross CT, Canteras NS (2012) The many paths to fear. Nat Rev Neurosci 13:651–658. doi: 10.1038/nrn3301

Hensman R, Guimaraes FS, Wang M, Deakin JFW (1991) Effects of ritanserin on aversive classical conditioning in humans. Psychopharmacology (Berl) 104:220–224. doi: 10.1007/BF02244182

Hindi Attar C, Finckh B, Büchel C (2012) The Influence of Serotonin on Fear Learning. PLoS One 7:e42397. doi: 10.1371/journal.pone.0042397

Holt DJ, Coombs G, Zeidan M a, et al (2012) Failure of neural responses to safety cues in schizophrenia. Arch Gen Psychiatry 69:893–903. doi: 10.1001/archgenpsychiatry.2011.2310

Homan P, Levy I, Feltham E, et al (2019) Neural computations of threat in the aftermath of combat trauma. Nat Neurosci 22:470–476. doi: 10.1038/s41593-018-0315-x

Hornung J (2010) The neuroanatomy of the serotonergic system. In: Muller CP, Jacobs BL (eds) Handbook of the Behavioral Neurobiology of Serotonin, 1st Ed. Elsevier, London, pp 51–64

Isosaka T, Matsuo T, Yamaguchi T, et al (2015) Htr2a-Expressing Cells in the Central Amygdala Control the Hierarchy between Innate and Learned Fear. Cell 163:1153–1164. doi: 10.1016/j.cell.2015.10.047

Kim MJ, Loucks R a., Palmer AL, et al (2011) The structural and functional connectivity of the amygdala: From normal emotion to pathological anxiety. Behav Brain Res 223:403–410. doi: 10.1016/j.bbr.2011.04.025

Lapiz-Bluhm MDS, Soto-Piña AE, Hensler JG, Morilak DA (2009) Chronic intermittent cold stress and serotonin depletion induce deficits of reversal learning in an attentional set-shifting test in rats. Psychopharmacology (Berl) 202:329–341. doi: 10.1007/s00213-008-1224-6

LeDoux JE (2000) Emotion Circuits in the Brain. Annu Rev Neurosci 23:155–184. doi: 10.1176/foc.7.2.foc274

Marin MF, Zsido RG, Song H, et al (2017) Skin conductance responses and neural activations during fear conditioning and extinction recall across anxiety disorders. JAMA Psychiatry 74:622–631. doi: 10.1001/jamapsychiatry.2017.0329

Matias S, Lottem E, Dugué GP, Mainen ZF (2017) Activity patterns of serotonin neurons underlying cognitive flexibility. Elife 6:1–24. doi: 10.7554/eLife.20552

Milad MR, Furtak SC, Greenberg JL, et al (2013) Deficits in conditioned fear extinction in obsessive-compulsive disorder and neurobiological changes in the fear circuit. JAMA Psychiatry 70:608–618. doi: 10.1001/jamapsychiatry.2013.914

Milad MR, Pitman RK, Ellis CB, et al (2009) Neurobiological Basis of Failure to Recall Extinction Memory in Posttraumatic Stress Disorder. Biol Psychiatry 66:1075–1082. doi: 10.1016/j.biopsych.2009.06.026

Murphy FC, Smith KA, Cowen PJ, et al (2002) The effects of tryptophan depletion on cognitive and affective processing in healthy volunteers. Psychopharmacology (Berl) 163:42–53. doi: 10.1007/s00213-002-1128-9

Nishizawa S, Benkelfat C, Young SN, et al (1997) Differences between males and females in rates of serotonin synthesis in human brain. Proc Natl Acad Sci 94:5308–5313. doi: 10.1073/pnas.94.10.5308

Öhman A (2009) Of snakes and faces: An evolutionary perspective on the psychology of fear. Scand J Psychol 50:543–552. doi: 10.1111/j.1467-9450.2009.00784.x

Park SB, Coull JT, McShane RH, et al (1994) Tryptophan depletion in normal volunteers produces selective impairments in learning and memory. Neuropharmacology 33:575–588. doi: 10.1016/0028-3908(94)90089-2

Passamonti L, Crockett MJ, Apergis-Schoute AM, et al (2012) Effects of acute tryptophan depletion on prefrontal-amygdala connectivity while viewing facial signals of aggression. Biol Psychiatry 71:36–43. doi: 10.1016/j.biopsych.2011.07.033

Patton, J.H., Stanford, M.S., Barratt E. (1995) Factor structure of the Barratt impulsiveness scale. J Clin Psychol 51:768–774

Phillips BU, Robbins TW (2020) The role of central serotonin in impulsivity, compulsivity and decision-making: comparative studies in experimental animals and humans. In: Muller CP, Cunningham KA (eds) Handbook of the Neurobiology of Serotonin, 2nd Ed. American Press, pp 531–548

Pruessner JC (2004) Dopamine Release in Response to a Psychological Stress in Humans and Its Relationship to Early Life Maternal Care: A Positron Emission Tomography Study Using [11C]Raclopride. J Neurosci 24:2825–2831. doi: 10.1523/jneurosci.3422-03.2004

Quednow B, Geyer M, Halberstadt A (2020) Serotonin and schizophrenia. In: Muller C, Cunningham KA (eds) Handbook of the Behavioral Neurobiology of Serotonin, 2nd Ed. Academic Press, pp 711–744

Raio CM, Hartley CA, Orederu TA, et al (2017) Stress attenuates the flexible updating of aversive value. Proc Natl Acad Sci 114:11241–11246. doi: 10.1073/pnas.1702565114

Remijnse PL, Nielen MMA, Van Balkom AJLM, et al (2006) Reduced orbitofrontal-striatal activity on a reversal learning task in obsessive-compulsive disorder. Arch Gen Psychiatry 63:1225–1236. doi: 10.1001/archpsyc.63.11.1225

Robbins TW, Vaghi MM, Banca P (2019) Obsessive-Compulsive Disorder: Puzzles and Prospects. Neuron 102:27–47. doi: 10.1016/j.neuron.2019.01.046

Robinson OJ, Overstreet C, Allen PS, et al (2012) Acute tryptophan depletion increases translational indices of anxiety but not fear: serotonergic modulation of the bed nucleus of the stria terminalis? Neuropsychopharmacology 37:1963–71. doi: 10.1038/npp.2012.43

Rogers RD, Everitt BJ, Baldacchino A, et al (1999) Dissociable deficits in the decision-making cognition of chronic amphetamine abusers, opiate abusers, patients with focal damage to prefrontal cortex, and tryptophan-depleted normal volunteersEvidence for monoaminergic mechanisms. Neuropsychopharmacology 20:322–339. doi: 10.1016/S0893-133X(98)00091-8

Schiller D, Levy I, Niv Y, et al (2008) From Fear to Safety and Back: Reversal of Fear in the Human Brain. J Neurosci 28:11517–11525. doi: 10.1523/JNEUROSCI.2265-08.2008

Schiller D, Kanen JW, LeDoux JE, et al (2013) Extinction during reconsolidation of threat memory diminishes prefrontal cortex involvement. Proc Natl Acad Sci U S A 110:20040–5. doi: 10.1073/pnas.1320322110

Seo C, Guru A, Jin M, et al (2019) Intense threat switches dorsal raphe serotonin neurons to a paradoxical operational mode. Science (80-) 363:539–542. doi: 10.1126/science.aau8722

Seymour B, Daw ND, Roiser JP, et al (2012) Serotonin Selectively Modulates Reward Value in Human Decision-Making. J Neurosci 32:5833–5842. doi: 10.1523/JNEUROSCI.0053-12.2012

Sheehan D, Lecrubier Y, Sheehan K, et al (1998) The Mini-International Neuropsychiatric Interview (MINI): the development and validation of a structured diagnostic psychiatric interview for DSM-IV and ICD-10. J Clin Psychiatry

Skandali N, Rowe JB, Voon V, et al (2018) Dissociable effects of acute SSRI (escitalopram) on executive, learning and emotional functions in healthy humans. Neuropsychopharmacology 43:2645–2651. doi: 10.1038/s41386-018-0229-z

Stahl SM (2013) Stahl’s Essential Psychopharmacology: Neuroscientific Basis and Practical Applications, 4th Ed. Cambridge University Press, Cambridge, UK

Talbot PS, Watson DR, Barrett SL, Cooper SJ (2006) Rapid tryptophan depletion improves decision-making cognition in healthy humans without affecting reversal learning or set shifting. Neuropsychopharmacology 31:1519–1525. doi: 10.1038/sj.npp.1300980

Terburg D, Morgan BE, Montoya ER, et al (2012) Hypervigilance for fear after basolateral amygdala damage in humans. Transl Psychiatry 2:. doi: 10.1038/tp.2012.46

van Donkelaar EL, Blokland A, Ferrington L, et al (2011) Mechanism of acute tryptophan depletion: Is it only serotonin. Mol Psychiatry 16:695–713. doi: 10.1038/mp.2011.9

Walker SC, Robbins TW, Roberts a. C (2009) Differential contributions of dopamine and serotonin to orbitofrontal cortex function in the marmoset. Cereb Cortex 19:889–898. doi: 10.1093/cercor/bhn136

Waltz JA, Gold JM (2007) Probabilistic reversal learning impairments in schizophrenia: Further evidence of orbitofrontal dysfunction. Schizophr Res 93:296–303. doi: 10.1016/j.schres.2007.03.010

Watson D, Clark LA, Tellegen A (1988) Development and validation of brief measures of positive and negative affect: the PANAS scales. J Pers Soc Psychol 54:1063–1070

Winstanley CA, Dalley JW, Theobald DEH, Robbins TW (2004) Fractioning impulsivity: Contrasting effects of central 5-HT depletion on different measures of impulsive behaviour. Neuropsychopharmacology 29:1331–1343. doi: 10.1038/sj.npp.1300434

Worbe Y, Savulich G, Voon V, et al (2014) Serotonin depletion induces “waiting impulsivity” on the human four-choice serial reaction time task: cross-species translational significance. Neuropsychopharmacology 39:1519–26. doi: 10.1038/npp.2013.351

Zangrossi H, Del Ben CM, Graeff FG, Guimarães FS (2020) Serotonin in Panic and Anxiety Disorders. In: Muller C, Cunningham KA (eds) Handbook of the Behavioral Neurobiology of Serotonin, 2nd Ed. Academic Press, pp 611–634

